# Estimating the sensorimotor components of cybersickness

**DOI:** 10.1101/318758

**Authors:** Séamas Weech, Jessy Parokaran Varghese, Michael Barnett-Cowan

**Author notes:** **Corresponding author:** Séamas Weech, Department of Kinesiology, University of Waterloo, Waterloo, Ontario, Canada.

## Abstract

The user base of the virtual reality (VR) medium is growing, and many of these users will experience cybersickness. Accounting for the vast inter-individual variability in cybersickness forms a pivotal step in solving the issue. Most studies of cybersickness focus on a single factor (e.g., balance, sex, vection), while other contributors are overlooked. Here, we characterize the complex relationship between cybersickness and several indices of sensorimotor processing. In a single session, we conducted a battery of tests of balance control, vection responses, and vestibular sensitivity to self-motion. A principal components regression model, primarily composed of balance control measures during vection, significantly predicted 37% of the variability in cybersickness measures. We observed strong, inverse associations between measures of sway and cybersickness. The results reiterate that the relationship between balance control and cybersickness is anything but straightforward. We discuss other factors that may account for the remaining variance in cybersickness.

## 1. Introduction

Virtual reality (VR) technology allows a user to experience a simulated environment through an array of sensory stimulation apparatuses (Hale & Stanney, 2014). Such arrays typically consist of electronic visual displays and sound devices, which can be updated in real-time based on the input of manual controllers, inertial motion units, and (depending on the hardware) eye tracking. Costs for these components have fallen in recent years, leading to the rapid adoption of VR hardware by enthusiasts. While the technology has a wealth of potential in a variety of settings, such as industrial skills training, consumer entertainment, and clinical rehabilitation, the sickness and discomfort experienced by many users of VR limit further adoption (Biocca, 1992; Keshavarz & Hecht, 2011; Kim et al., 2005). Sickness during VR exposure, termed cybersickness (CS), has been studied in some detail in recent years. The phenomenon is related to several maladies under the general term ‘motion sickness’, including car-sickness or sea-sickness (Riccio & Stoffregen, 1991), visually-induced motion sickness (Graybiel & Miller, 1974; Keshavarz et al., 2015), and simulator sickness (Kennedy et al., 1993), which each result from exposure to different manners of novel sensory environments. Symptoms are wide-ranging, including nausea, skin pallour, headache, disorientation, ocular discomfort, and in extreme cases, vomiting (LaViola Jr, 2000). Although there are several hypotheses about the causes of these symptoms (Golding & Gresty, 2005; Johnson et al., 1999; Keshavarz et al., 2015; Oman, 1990; Riccio & Stoffregen, 1991), existing theories have yet to offer techniques that prevent the occurrence of CS. This problem requires a solution if we are to benefit from the potential impact of VR technology on society. The fact that large individual differences exist in terms of CS susceptibility suggests that some factors that differ between susceptible and non-susceptible users can be identified and used to guide the development of CS prevention methods. Existing literature has identified several factors that may explain individual heterogeneity in CS, but most studies focus on one rather than multiple contributing factors. Here, we first provide an overview of the literature with a focus on highlighting evidence for a multifactorial causal structure for CS. We then describe an experiment in which we collected several measures of sensorimotor processing (e.g., balance control, self-motion sensitivity) before participants were exposed to VR, and used these measures to construct a multiple regression model with the aim of predicting the severity of CS.

### 1.1. Balance control

Recent research has suggested that individual differences in CS are related to balance control and self-motion perception (Dennison & D’Zmura, 2017; Keshavarz et al., 2015; Riccio & Stoffregen, 1991; Sadiq et al., 2017). Perceiving and controlling self-motion requires the integration of multisensory cues (e.g., vision, audition, proprioception, & vestibular) to derive knowledge about the state of the body in space. VR exposure can lead to a ‘ sensory rearrangement’ (Reason & Brand, 1975; Welch, 2002) where the learned relationships between sensory modalities are modified; for example, in VR, small but critical delays between sensory feedback across modalities can affect the perception and control of the temporal evolution of an action (Biocca et al., 2001). As well, visual and vestibular cues that convey information about the state of the head-on-body are frequently incongruent in VR, which may pose a challenge for the maintenance of stable postural control. The ‘postural instability theory’ of sickness was formalised by Riccio and Stoffregen (1991; also see Chardonnet et al., 2017; Stoffregen & Riccio, 1991; Stoffregen & Smart, 1998; Takada et al., 2007), who suggested that motion sickness emerges as a consequence of postural instability resulting from unfamiliar environmental conditions. In support of this theory lies evidence that motion sickness produced by simple optic flow stimuli is predicted by the temporal dynamics of postural activity (Palmisano et al., 2018). Others have shown that the area of postural sway tends to increase when participants experience CS (Chardonnet et al., 2017). Interestingly, while several of these studies have documented a positive correlation between postural sway and measures of CS, other studies have found no evidence of this link (Dennison & D’Zmura, 2018), while yet others have observed that individuals who experience stronger CS tend to demonstrate decreased postural sway (Dennison & D’Zmura, 2017; Sadiq et al., 2017). These authors concluded that participants who experience CS may desire to remain stationary in order to avoid increased exposure to decoupled sensory streams in VR, resulting in reduced postural sway for individuals who are highly susceptible to CS (this phenomenon has been termed ‘VR lock’). Taken together, the results suggest that there is a relationship between CS and balance control, but also a need for further examination, particularly to characterize how other factors might modulate the relationship.

### 1.2. Visual motion perception and vection

Consistent with the theory that CS arises due to stresses imposed on the sensorimotor control system by VR, recent evidence shows that individual variability in visual motion sensitivity may explain part of the heterogeneity in CS susceptibility rates (Allen et al., 2016). Individuals with a greater sensitivity to 3D visual motion are more likely to opt for early termination during exposure to nauseogenic VR conditions, and tend to experience higher levels of discomfort. Participants with greater visual sensitivity may have been more likely to detect the cue-conflicts that occur during VR use, such as the visual-vestibular mismatch produced when self-motion is simulated with optic flow in VR. Notably, the stereoscopic 3D motion stimulus presented by Allen and colleagues (2016) is highly relevant to the flow parsing process that underpins self-motion perception, and as such, heterogeneity in susceptibility to visual self-motion illusions may explain why self-motion in VR results in sickness in some but not others (Keshavarz et al., 2015). It is possible that measures of vection (visually-induced perception of self-motion; Dichgans & Brandt, 1978) during visually-simulated self-motion could reveal correlated differences in CS and visual self-motion sensitivity. Although some have characterized the effect of vection-inducing stimuli on vection ratings and postural sway (Berthoz et al., 1979; Palmisano et al., 2014) and shown that strong vection predicts high simulator and motion sickness (Hettinger et al., 1990; Hettinger & Riccio, 1992; for a review, see Keshavarz et al., 2015), others have reported a negative relationship between vection and CS (Palmisano et al., 2017), or no relationship (Palmisano et al., 2018). The agreed consensus appears to be that the relationship is highly complex and requires further examination (Keshavarz et al., 2015; Palmisano et al., 2017) with more advanced analysis techniques.

### 1.3. Vestibular sensitivity

It is likely that differences in vestibulo-inertial perception play a key role in the variability observed in CS across individuals. Clinical research on vestibular dysufunction patients has been important in this context, showing that individuals with vestibular labyrinth lesions do not exhibit sickness when exposed to a rotating visual field stimulus (Cheung et al., 1991, 1989; Johnson et al., 1999). When healthy participants are exposed to the same stimulus, symptoms that mirror those of CS result. The vestibular sense also plays a crucial role in the maintenance of balance control through detecting fluctuations in the inherently unstable human body (Peterka, 2002). Given the relationship between postural instability and CS proposed by Riccio and Stoffregen (1991) and supported by others (e.g., Chardonnet et al., 2017; Takada et al., 2007), it is clear that vestibular sensitivity to self-motion is likely to modulate CS to some degree. Further support for this point arises from evidence of a strong comorbidity between vestibular migraine and sickness susceptibility (Money & Cheung, 1983). Susceptibility to other forms of sickness (e.g., motion/space-flight sickness) is related to individual differences in vestibular functioning (Diamond & Markham, 1991; Hoffer et al., 2003; Quarck et al., 2000). For instance, Diamond and Markham (1991) found differences in spontaneous ocular torsion between astronauts who experienced sickness during space-flight compared to those who did not. Recent efforts to reduce visual-vestibular cue mismatch in VR support a partial vestibular basis for CS: Both galvanic vestibular stimulation (Cevette et al., 2012; Gálvez-García et al., 2015; Reed-Jones et al., 2007) and bone vibration applied near the vestibular organs (Weech et al., 2018) reduce the level of CS experienced during VR use. There is also a striking similarity between CS symptoms and the symptoms of vestibular labyrinthectomy—although the former are less severe than the latter— with the effects of labyrinthectomy including “nausea and vomiting… excessive perspiration… The face is pallid and the skin is clammy. The patient. resists any head movement for fear that any alteration will increase the vertigo and bring on a spell of severe nausea and vomiting” (Gernandt, 1959, p.562). The results of a study of vestibular migraine patients suggested an increased ‘sensitivity’ to vestibular cues in such patients, such that undergoing a rapid spin was likely to result in motion sickness for such patients (Kuritzky et al., 1981). These authors took this as evidence that the increased motion sickness suffered by vestibular migraine patients was attributable to lower vestibular thresholds. The vestibular system is also strongly implicated in the magnitude and latency of vection onset (Weech & Troje, 2017; Wong & Frost, 1981), which has a complex relationship with CS as discussed above. Despite the strong evidence that motion sickness appears to be partially attributable to high vestibular sensitivity, there are no studies to our knowledge that have investigated the relationship between vestibular self-motion sensitivity and CS.

### 1.4. Additional factors

Symptoms of CS and other forms of motion sickness are correlated with heightened autonomic nervous system activity (Harm, 2002; Ohyama et al., 2007). Common measures of CS emphasize qualitative experiences that indicate abnormal physiological functioning, such as excess sweating, burping, and stomach awareness (Kennedy et al., 1993). Marked differences in hormonal secretion (e.g., adrenocorticotropin and growth hormone) have been observed between individuals who are susceptible to motion sickness (Kohl, 1985; Eversmann et al., 1978). Although it is unclear if these physiological correlates are causally related to the qualitative experience of CS, it is thought that physiological characteristics of CS may represent the process of sensory conflict reduction (Harm, 2002; Welch, 2002). Physiological measurements such as electroencephalography (EEG) and electrocardiography (ECG) demonstrate predictive validity for self-reported CS scores (Dennison et al., 2016; Kim et al., 2005). However, these measures are typically obtained during VR exposure when CS symptoms have already emerged. The practical utility of predicting CS from on-line measurements is limited by the fact that CS symptoms can emerge quickly and can be long-lasting, even upon exiting the nauseogenic conditions (Bos, 2011; Kennedy et al., 2000; Regan, 1995). Although it may be possible to gather physiological data prior to VR exposure and use those to predict the emergence of CS, we are not aware of any studies that have used such an approach.

### 1.5. Approach of the current study

The studies discussed here provide a wealth of evidence about the possible causes of heterogeneity in CS. Principally, these factors include balance control (Chardonnet et al., 2017; Riccio & Stoffregen, 1991; Takada et al., 2007), vestibular motion sensitivity (Cheung et al., 1991; Diamond & Markham, 1991; Hoffer et al., 2003), and visual motion/self-motion perception (Allen et al., 2016; Keshavarz et al., 2015). However, no studies have assessed the relative impact of each factor. Quantifying the role of each requires the assessment of responses in several behavioural tasks. These data, which are illustrative of the sensorimotor control system of a participant, may be used to construct a model that predicts the likelihood that the individual will experience cybersickness—without exposing the individual to the unpleasantness of such an experience.

The purpose of the present study was to characterise the degree to which CS susceptibility is attributed to individual differences in balance control, and self-motion perception from visual and vestibular cues. We predicted that balance control would account for a large proportion of variability in CS scores, due to research suggesting that poor balance control precedes CS (Riccio & Stoffregen, 1991). We also expected that susceptibility to vection would predict CS based on the association between vection and visually-induced motion sickness (Keshavarz et al., 2015). Finally, motivated by literature that shows increased vestibular sensitivity predicts high susceptibility to motion sickness (Diamond & Markham, 1991; Money & Cheung, 1983), we expected that high vestibular self-motion sensitivity would be predictive of high CS scores. However, our main goal was to establish a multifactorial statistical model for predicting CS based on a combination of these factors.

## 2. Methods

### 2.1. Participants

#### 2.1.1. Recruitment

Participants were recruited using mailing lists and posters on the University of Waterloo campus and were remunerated $10 per hour of participation. Participants were all naïve to the purpose of the experiment. This study was carried out in accordance with the recommendations of Canada’s Tri-Council Policy Statement: Ethical Conduct for Research Involving Humans (TCPS2) by the University of Waterloo’s Human Research Ethics Committee with written informed consent from all subjects. All subjects gave written informed consent in accordance with the Declaration of Helsinki. The protocol was approved by the University of Waterloo’s Human Research Ethics Committee.

#### 2.1.2. Demographics

Thirty undergraduate and graduate students participated in the study (12 were male, *M* age = 22.87, *SD* = 3.94, range = 18-30). All participants had normal or corrected-to-normal vision, and reported having no musculoskeletal, neurological, or balance disorders.

Participants were invited to optionally record their ethnicity, and 23 of the 30 participants chose to do so. Of those who reported ethnicity, 10 reported ‘Asian’ and 13 reported ‘European’. In addition participants were invited to record their daily activity level (low/moderate/high). In total, seven reported ‘low’ daily activity, eighteen reported ‘moderate’ activity, and the remaining five reported ‘high’ daily activity.

### 2.2. General Procedure

The general procedure of the experiment is described here, and further details are provided for each task in a separate section below (2.3. Specific procedures).

Before commencing the experiment, participants were introduced to the purpose of the study through the letter of information and consent. Participants then completed a questionnaire to assess the incidence of motion sickness in their daily activities and in childhood (Motion Sickness Susceptibility Questionnaire, MSSQ; Golding, 1998) and reported demographic information. Following this, participants were asked to optionally supply a saliva sample to be processed for allele group identification.

In the second part of the study participants completed balance control and self-motion tasks in a block design that was counterbalanced across participants. In the balance control tasks, participants were guided through the process of assessing their balance using force-plates in five different sensory conditions (outlined in detail below, see 2.3.1. Balance control task). In one of the balance tasks, we presented participants with a radial optic flow stimulus and assessed their visually-induced sway as an index of vection susceptibility. In the vestibular self-motion sensitivity task, participants were passively rotated in yaw while seated on a motion base and were asked to report their direction of rotation (left/right).

In the third part of the study, participants completed two VR tasks and reported the level of post-exposure CS. The VR tasks were administered with a predetermined order that was counterbalanced across participants. This part of the study was completed last in the experimental sequence in order to avoid any possible effects of sickness on performance in the other tasks. The total duration of the experiment was approximately 150 min.

### 2.3. Specific procedures

#### 2.3.1. Balance control task

The balance control task comprised five sensory conditions in a block design where measures of postural sway were collected using two force-plates (4060-05, Bertec Corporation, Columbus, OH, USA) arranged in a side-by-side configuration, separated by approximately 1 cm. Prior to data collection, participants were asked to stand unshoed in a standardized foot position (approximately shoulder width stance with toes rotated laterally by 14 degrees; McIlroy & Maki, 1997; see Figure 1) with each foot on one of the two force-plates. The outline of the feet was traced with markings to ensure consistent orientation of the feet across participants. After establishing the initial stance position, the participants were asked to stand quietly for 30 sec with hands at their side in five sensory conditions that manipulated their visual or somatosensory inputs. Figure 1 depicts the conditions of the balance control task. The five sensory conditions were standing with eyes open (eyes-open standard, EOS), standing with eyes closed (eyes-closed standard, ECS), standing on foam with eyes open (eyes-open foam, EOF) or standing on foam with eyes closed (eyes-closed foam, ECF), and standing while observing a radial optic flow stimulus that induced vection (V; see 2.3.2. Vection task). In eyes-open (EO) conditions, the participants were asked to fixate on a cross (5 cm) placed at eye level on a wall 2.74 m in front of the participant (visual angle of the cross was approximately 1 x 1 deg). After each trial participants were required to take a 10 sec break. Trials in this task were blocked by sensory condition and administered in a predetermined randomized order. If the participant intentionally moved or stepped off the force-plates during a trial, the trial was recollected. The task lasted approximately 45 min including setup of the apparatus and instructions.

**Figure 1.**
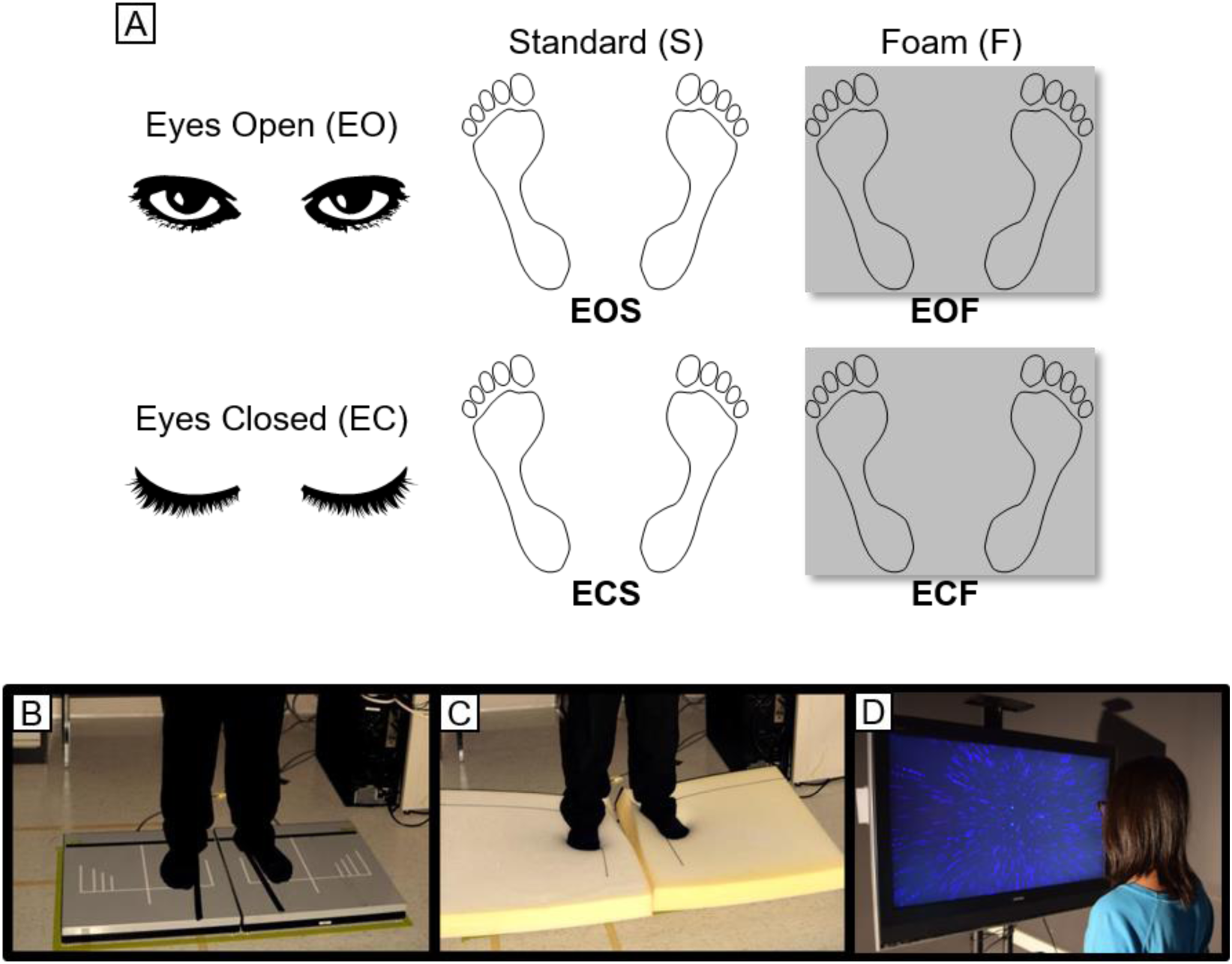
(A) Design of balance tasks: EOS, EOF, ECS, and ECF. (B) Depiction of standard stance on force-plates (standard conditions). (C) Standing on foam that covered the force-plates (foam conditions). (D) Observing a radial optic-flow stimulus on a monitor (condition V).

Vertical ground reaction force (Fz) and moments of force (x and y planes) from the force plates were recorded over a 30 sec period for 8 trial repetitions for each sensory condition. The force plate data was amplified online using internal digital preamplifier and sampled at a rate of 1000 Hz and stored for offline analysis. The force plates were calibrated prior to data collection for each participant. The force plate data were acquired using a custom-built LabVIEW program (National Instruments, Austin, TX, USA). Our choice of trial-duration (30 sec) was motivated by evidence that this duration provides the optimum test-retest reliability (Clair & Riach, 1996). In addition, 30 sec stance is a common standard for postural sway measurement in adults and clinical population because longer durations (one min or more) may be too lengthy for a patient (Duarte & Freitas, 2010).

#### 2.3.2. Vection task

As part of the balance control section participants observed a vection stimulus (condition V) while balance measures and verbal reports were collected. Participants observed a radially expanding optic flow stimulus (Figure 2) that consisted of a cloud of 1000 randomly-positioned dots (0.25 deg visual angle; blue dots on a black background; a video depiction is provided in Movie 1: https://www.dropbox.com/s/akzht7r80cda2u1/vection_stimulus_movie.avi?dl=0). The movement of the dots towards the observer was intended to give rise to the impression of linear translation of the observer in the anterior-posterior axis. The dots contained linear perspective and relative size cues to depth. The visual stimulus included an oscillation component (0.5 Hz mediolateral frequency; 1 Hz ventrodorsal frequency) which is known to enhance the sense of vection (e.g., Apthorp & Palmisano, 2014; Palmisano & Kim, 2009).

**Figure 2.**
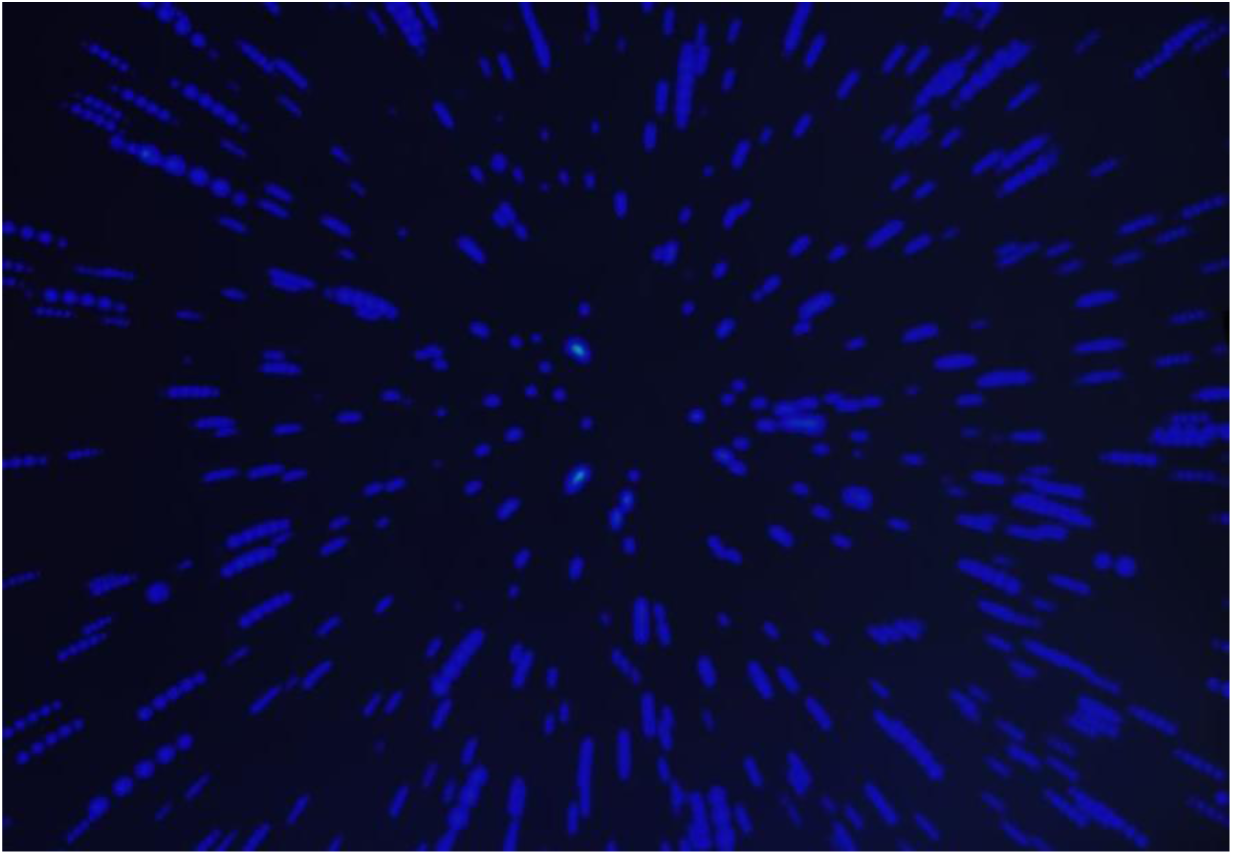
Long exposure of the vection stimulus (also see Movie 1).

The optic flow stimulus was presented on an LCD screen (76 × 133 cm) which was adjusted to eye height and positioned 53 cm ahead of the observer (visual angle was approximately 71 × 103 deg). Before the block of vection trials, the experimenter explained the feeling of vection (“You may feel the illusion that your own body is moving through space”). The investigator provided the example that vection can occur when looking out of a window at a moving vehicle. Participants were required to verbally confirm that they understood what was meant by vection. Each participant was shown an example of the vection stimulus and asked if they indicated vection. All participants except one reported vection (note, data from this participant were not excluded from analyses). Next, the experimenter instructed participants that they were required to indicate how strongly they felt vection after each trial on a scale of 0 to 10. The anchors provided were “0: No vection at all” to “10: The strongest possible feeling of vection”. Participants were positioned in front of the LCD display while they stood on the force plates. Before each trial began participants fixated on a central cross (~0.5 deg) on the LCD display that specified where gaze should be located during the trial. The fixation cross disappeared once the vection stimulus began. The vection stimulus was presented for 30 sec, during which balance control data were obtained from the force plates. Finally, the experimenter asked the participant to verbally report the strength of vection experienced during the trial on the ‘0 to 10’ scale.

#### 2.3.3. Vestibular self-motion sensitivity task

We measured vestibular sensitivity to self-motion in terms of the ability of the participant to estimate the direction of yaw rotation on a motion platform when visual, auditory, and proprioceptive cues were minimised.

Participants were seated on a racing chair (A4, Corbeau LLC, Sandy, UT, USA) that was mounted to a motion base (MB-E-6DOF/12/1000KG, Moog Inc., Elma, NY, USA; Figure 3) with a custom-built frame. A five-point harness was used to ensure the participant’s position was stable, and the participant’s head was secured in place with a helmet. The vertical and horizontal location of the helmet was adjusted by the researcher to comfortably fit the head of each participant and to ensure that the axis of rotation of the motion platform intersected with the centre of the participant’s head. Participants used a blindfold, earplugs, and were exposed to white-noise auditory masking with active noise-canceling headphones while seated on the platform. Foam padding was mounted to the surface of the chair and the platform beneath the feet of participants in order to reduce the potential influence of proprioceptive cues. Participants were required to wear long sleeves and nitrile gloves in order to avoid an influence of air resistance on direction judgments. For the same reason, a fan mounted to the platform was directed at the face of the participant throughout the task.

**Figure 3.**
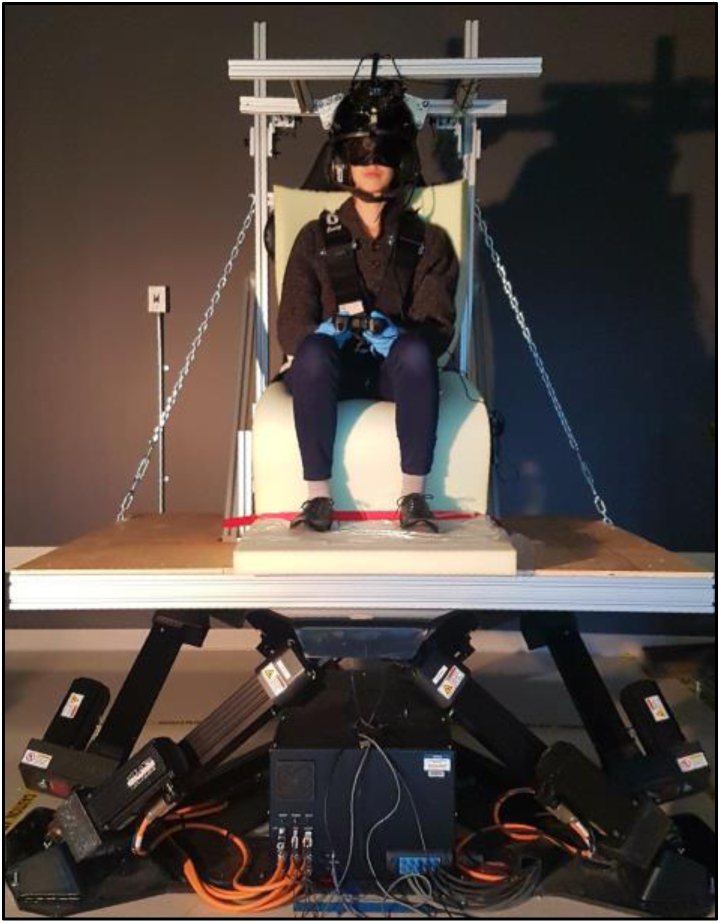
Apparatus used in the vestibular self-motion sensitivity task. Participants were seated on a cushioned seat mounted to a motion base and required to use a blindfold, earplugs, gloves, headphones, five point harness, and a helmet. In each trial the seat was rotated in yaw and the participant indicated the direction of rotation with a handheld gamepad.

Participants were rotated in yaw either left or right for 1 second in each trial according to raised cosine bell velocity profiles. The magnitude of rotation was selected trialwise from a list of 20 velocity profiles using a Bayesian adaptive staircase procedure, Psi-marginal (Kontsevich & Tyler, 1999). Peak velocity spanned a logarithmic space from 0.05-6.0 deg/sec. In each trial, a beep was played to signal movement onset (440 Hz, 250 ms), then the participant was rotated in yaw, after which a second beep was presented (880 Hz, 250 ms) to indicate that the participant should input their response (i.e., which direction they rotated: left or right). The response was input using a handheld gamepad (Logitech F310) where two face buttons indicated ‘left’ or ‘right’. The response value was used to update the estimate of the psychometric function in the Psi-marginal algorithm. A third beep was played after the response had been inputted (220 Hz, 250 ms). After the response was inputted, the motion base slowly rotated to its initial orientation in preparation for the next trial (7 sec; maximum possible velocity of the return movement was 0.85 deg/s which is sub-threshold at this frequency, Grabherr et al., 2008). A practice trial at the start of the task consisted of the same movement for each participant (2.1 deg/s).

The 150 total trials of this task were split into separate adaptive staircases for each direction (left/right) with 75 trials in each. Each trial lasted approximately 13 sec (0.25 sec beep, 1 sec movement, 0.25 sec beep, ~3 sec response, 7 sec return to centre of rotation, 1.5 sec pause). The task lasted approximately 45 min including setup of the apparatus and instructions.

#### 2.3.4. VR tasks

Participants were guided through the process of fitting a head mounted display (Rift CV1, Oculus VR, Menlo Park, CA, USA) and calibrating the device (interocular distance and position on face) before completing the VR tasks. Participants were asked to play two types of VR content which have been rated on the Oculus Store Comfort Rating system as either ‘intense’ or ‘comfortable’ (Figure 4). The two VR tasks were completed sequentially in a predetermined order which was counterbalanced across participants. Each task lasted 7 min in total.

**Figure 4.**
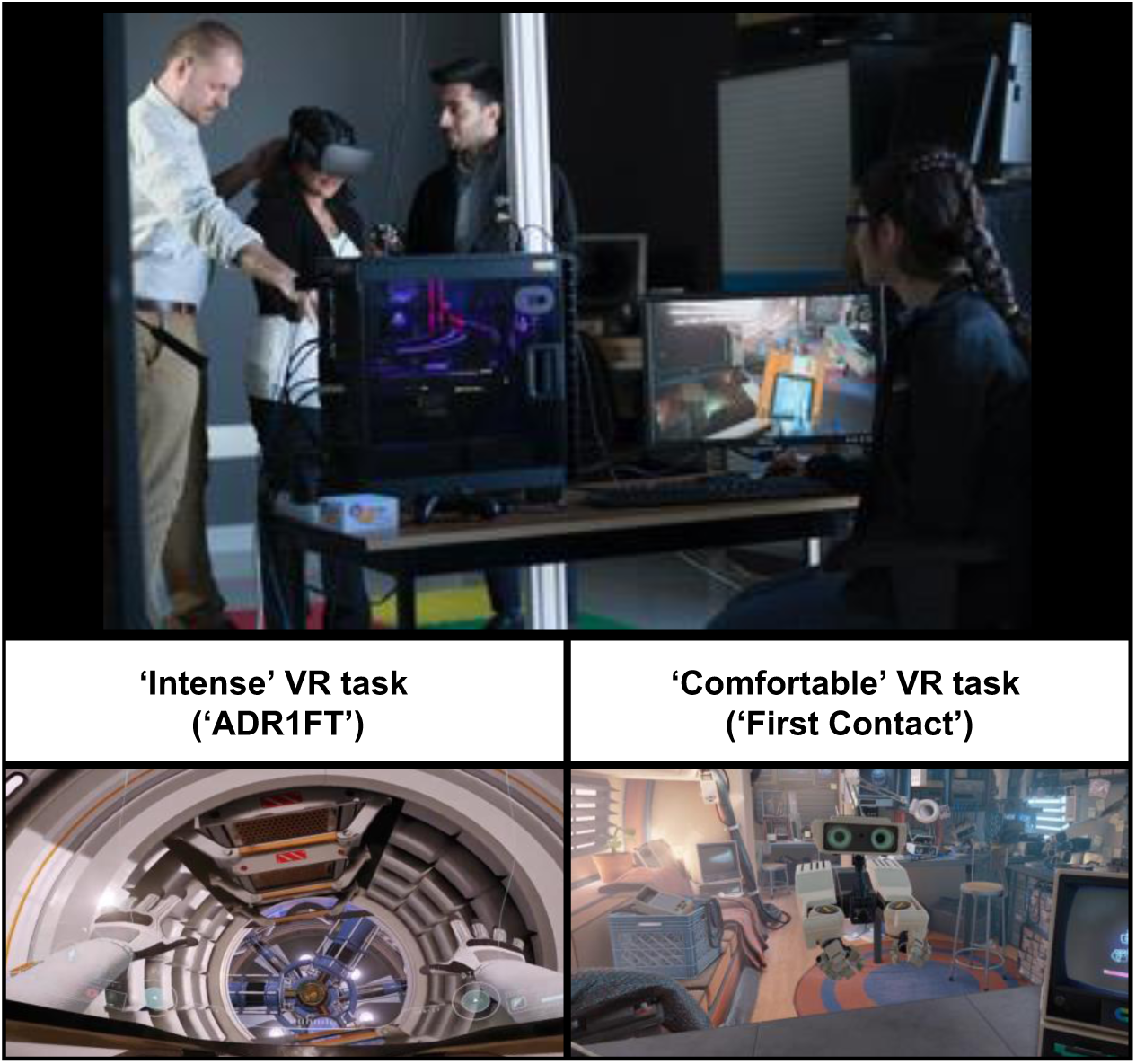
Top, a participant is introduced to the environment in one of the two VR tasks, which are depicted in the screenshots below.

The first VR task was ‘ADR1FT’, in which participants played the role of an astronaut freely exploring a simulated space station. The participant navigated through the environment using a handheld gamepad (Xbox One Wireless Controller, Microsoft, Redmond, WA, USA). This experience is rated as ‘intense’ on the Oculus Store (https://oculus.com/experiences/rift/905830242847405/).

The second VR task was ‘First Contact’, in which participants observed and communicated with a robot in a simulated environment that depicted the interior of a travel trailer. The participant was encouraged by the robot to perform simple actions such as picking up and throwing objects using six-degrees-of-freedom motion-tracked controllers (Touch Controllers, Oculus VR) that were held in the right and left hands. This experience is rated as ‘comfortable’ on the Oculus Store (https://oculus.com/experiences/rift/1217155751659625/).

After each VR task, participants completed the Simulator Sickness Questionnaire (SSQ; Kennedy et al., 1993), a checklist of 16 symptoms (e.g., nausea, headache, sweating) to be rated on a scale of ‘none’, ‘slight’, ‘moderate’ or ‘severe’. The VR tasks lasted approximately 30 min including setup of the apparatus and instructions.

### 2.4. Data Analysis

#### 2.4.1. Balance control data

Post-processing of forceplate data consisted of low pass filtering (6-Hz, dual-pass 2nd- order Butterworth filter), calculation of COP in anterior-posterior and medial-lateral positions, and extraction of COP parameter (sway path length) using a custom-made LabVIEW program. Sway path length is defined as the total length of the COP path in 30 sec, which is approximated by the sum of the distances between two consecutive points on the COP path (Hufschmidt et al., 1980; Prieto et al., 1996). The trialwise averages from all balance tasks were normally distributed (non-significant Kolmogorov-Smirnov tests, *Ds* ≤ 0.13, *ps* ≥ .64). In addition, we observed no significant outliers in any condition (one-sample Dixon outlier tests, *Qs* ≤ 0.34, *p*s ≥ .093).

Figure 5 depicts the total sway path length for each condition. To assess if balance control differed between the conditions we conducted a one-way repeated measures analysis of variance (ANOVA) with a conservative Greenhouse-Geisser correction. We found a significant difference between the conditions (F(3.04, 88.14) = 95.12, *p* < .001). Next we conducted planned paired-samples t-tests based on our expectation that closing the eyes, standing on foam, and experiencing vection would increase balance control variability. The vection condition resulted in significantly higher sway path length (*M* = .306 m, *SEM* = .011 m) than the EOS condition (*M* = .284 m, *SEM* = .010 m) which was our baseline comparison condition (*t*(29) = 2.87, *p* = .007). In addition, sway path length was lower in eyes open compared to eyes closed conditions, both standing on foam and in standard stance (*t*(29)s ≥ 6.87, *p*s < .001), and sway path length was lower in standard standing conditions compared to foam standing conditions in both eyes open and closed visual conditions (*t*(29)s ≥ 9.18, *p*s < .001).

**Figure 5.**
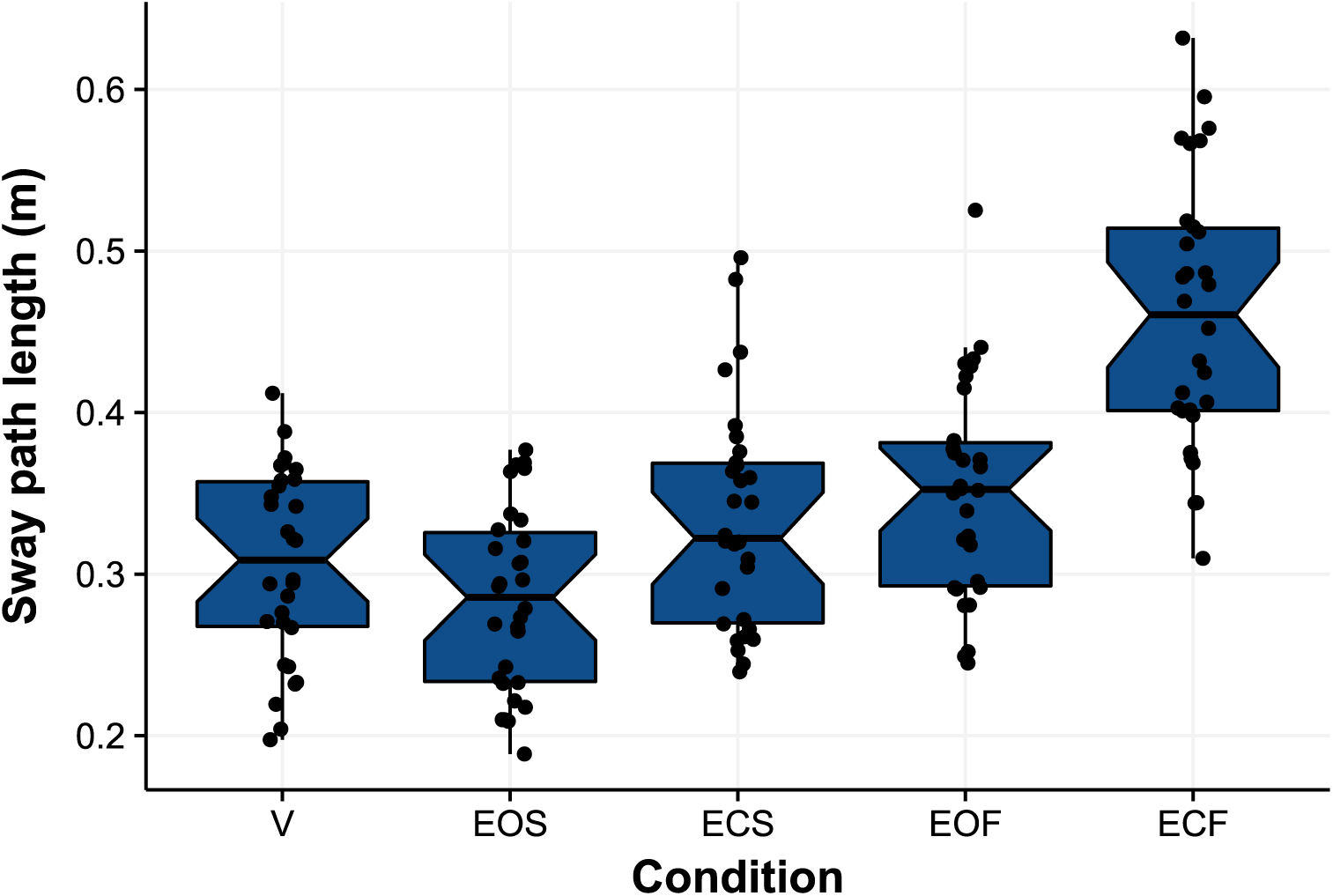
Notched box-plot depicting balance control measures for each condition. Thick horizontal lines are group averages. Black dots are participant averages. Note, while one datapoint for EOF was outside of 1.5 interquartile ranges, this was not a significant outlier (one-sample Dixon test, *Q* = 0.34, *p* = .093).

#### 2.4.2. Vection strength data

Participants’ estimates of the strength of vection experienced over eight trials were averaged to compute a single score for each participant. These data were normally distributed (non-significant Kolmogorov-Smirnov test, *D* = 0.13, *p* = .66). At the end of the block of vection trials we asked participants if they experienced any sickness during the vection trials, and received no affirmative responses.

Of the 30 participants in total, 29 reported feeling vection (strength ratings across all participants: *M* = 3.30, *SEM* = 0.34). Rather than discarding the data of the participant who reported no vection (e.g., Riecke & Feuereissen, 2012; Tarita-Nistor et al., 2008; Trutoiu et al., 2007), we retained their data for further analyses in order to avoid a possible sample bias.

#### 2.4.3. Vestibular self-motion estimation data

Direction discrimination reports (‘left’ or ‘right’) for each trial were used to update a standard algorithm for threshold estimation (Psi-marginal, Kontsevich & Tyler, 1999). Low threshold values indicate high sensitivity to self-motion and vice-versa. Since data obtained from the left and right staircases were not statistically different across participants (*p* > .05), we combined the left/right thresholds for each participant to obtain a single value depicting the 75% threshold for direction discrimination. Data for a single participant are depicted in Figure 6.

**Figure 6.**
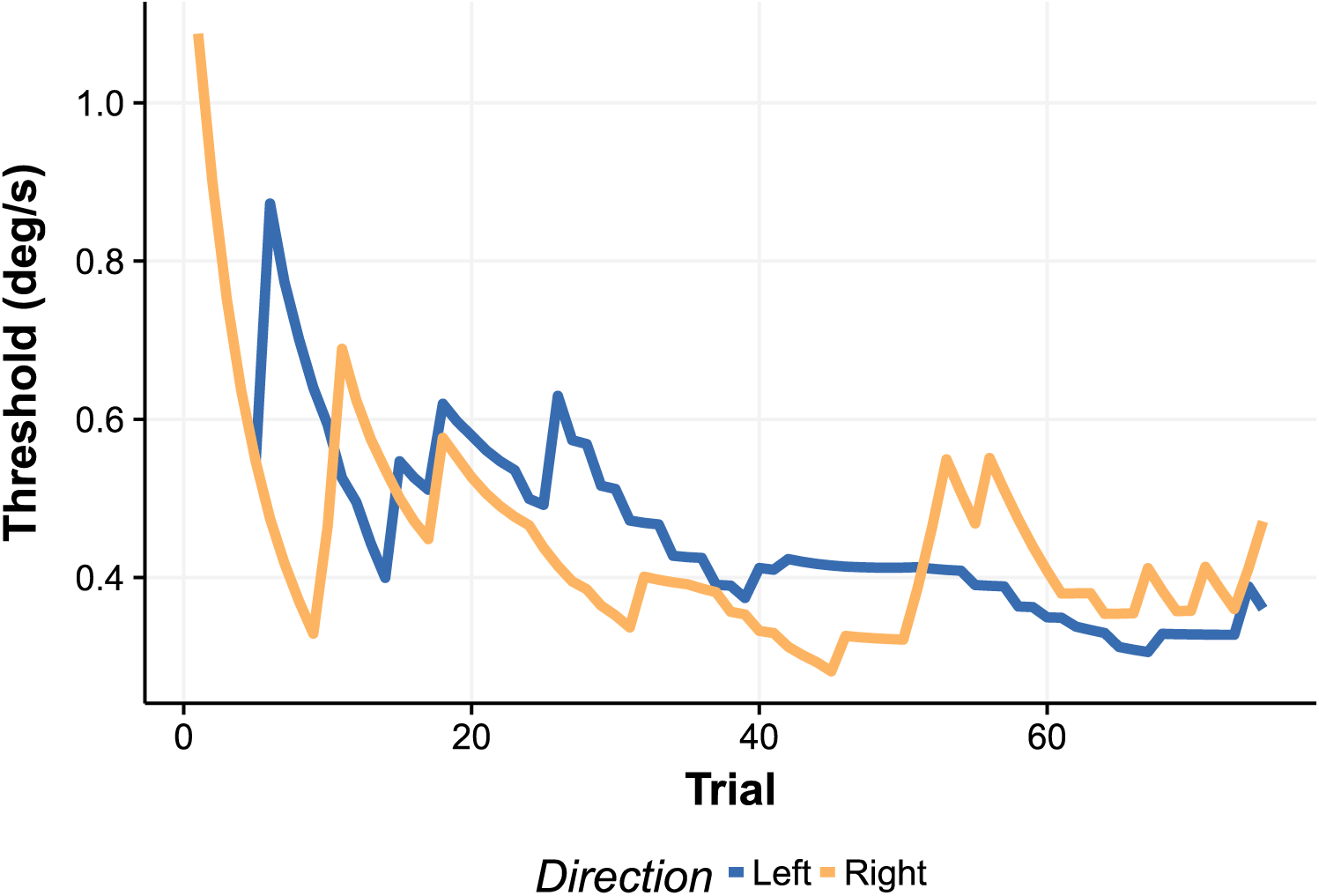
Threshold estimates for the left and right directions for a typical participant across 75 trials. The final value obtained from the staircase procedure averaged over the left and right directions (in this case approx. 0.4 deg/s) was taken as the best estimate of the threshold.

The vestibular threshold data obtained across participants were non-normally distributed (significant Kolmogorov-Smirnov test, *D* = 0.27, *p* = .02, see Figure 7) so we applied a square root transform to the data (non-significant Kolmogorov-Smirnov test, *D* = 0.21, *p* = .11) before subjecting the data to further analysis.

**Figure 7.**
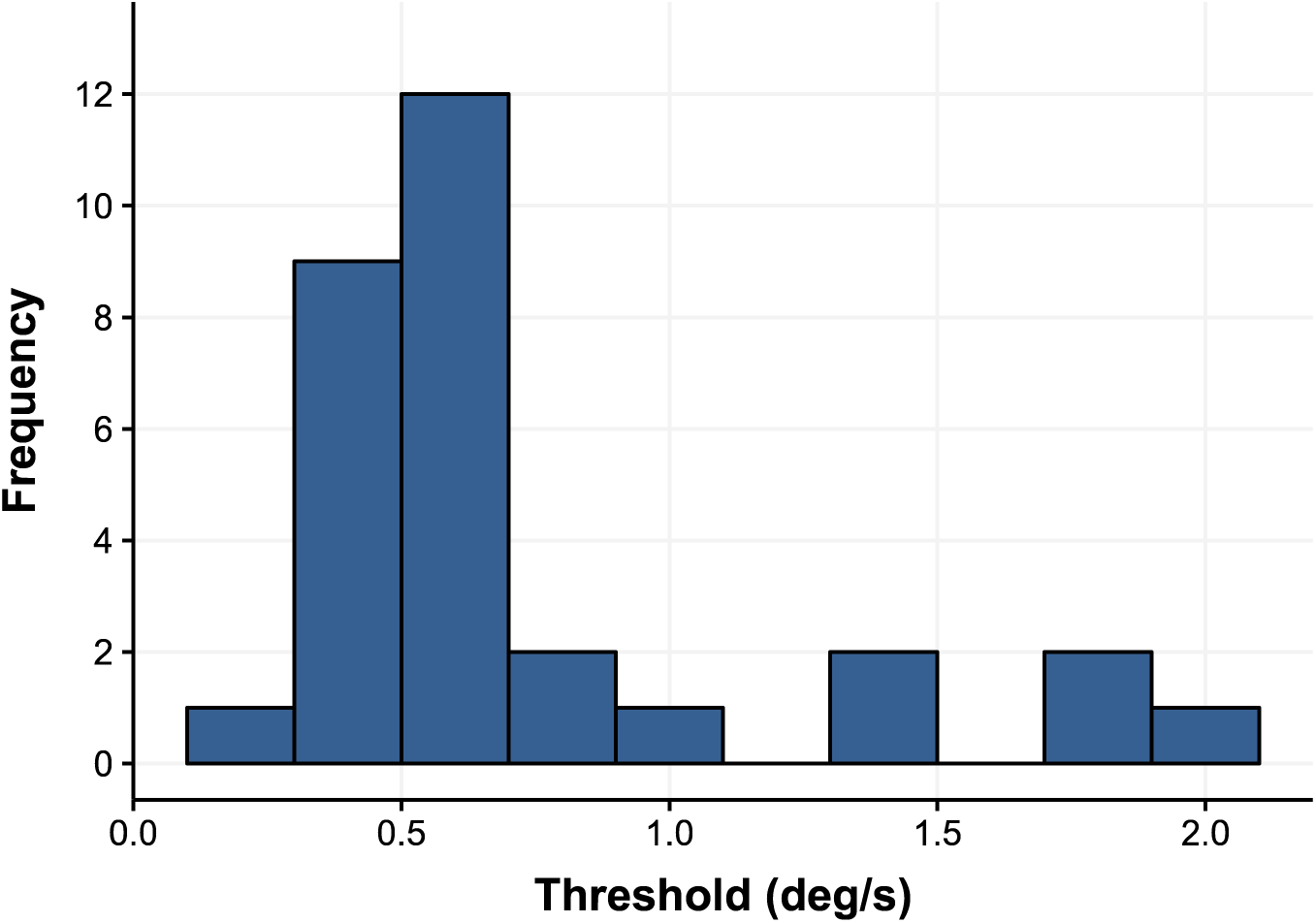
Histogram of vestibular threshold data (bin width = 0.2 deg/s).

#### 2.4.4. Cybersickness data

We used participants’ responses on the SSQ to compute total scores for each VR task according to the formula of Kennedy and colleagues (1993). To verify that we were accurate in characterising the two tasks as “intense” and “comfortable”, we compared the scores on each task and found a significant difference in sickness between the “intense” task (*M* = 37.90, *SEM* = 5.93) and the “comfortable” task (*M* = 10.22, *SEM* = 1.69), in agreement with our expectations (Welch’s *t*(33.69) = 4.49, *p* < .001).

We used the total scores on the SSQ for both VR tasks (“intense”: ADR1FT; “comfortable”: First Contact) to compute a difference score representing the effect of nauseogenic VR content on the participant’s comfort level, which we term ‘∆CS scores’. The ∆CS scores were normally distributed (non-significant Kolmogorov-Smirnov test, *D* = 0.18, *p* = .28). Although other research has characterised ‘cybersickness’ as SSQ scores on a single VR task to characterise sickness, we note that the ∆CS scores that we used here (i.e., difference in SSQs obtained after intense and comfortable VR content) were highly correlated with SSQ total scores for the “intense” VR content (*r*(28) = 0.95*, p* < .001). On the other hand, while the correlation between ∆CS scores and SSQ total scores for the “comfortable” VR content was also significant, the relationship was weaker (*r*(28) = 0.36, *p* = .049).

## 3. Results

### 3.1. Correlations and between-group effects

We conducted correlations to establish relationships between predictors and outcomes. A correlation plot of the data is depicted in Figure 8. First, as expected, we identified a significant positive correlation between ∆CS and past motion sickness susceptibility (MSSQ; *r*(28) = 0.36, *p* = .048), suggesting that individuals who often experience motion sickness in provocative situations (e.g., boats, fairground rides) were also more likely to experience CS in virtual environments.

**Figure 8.**
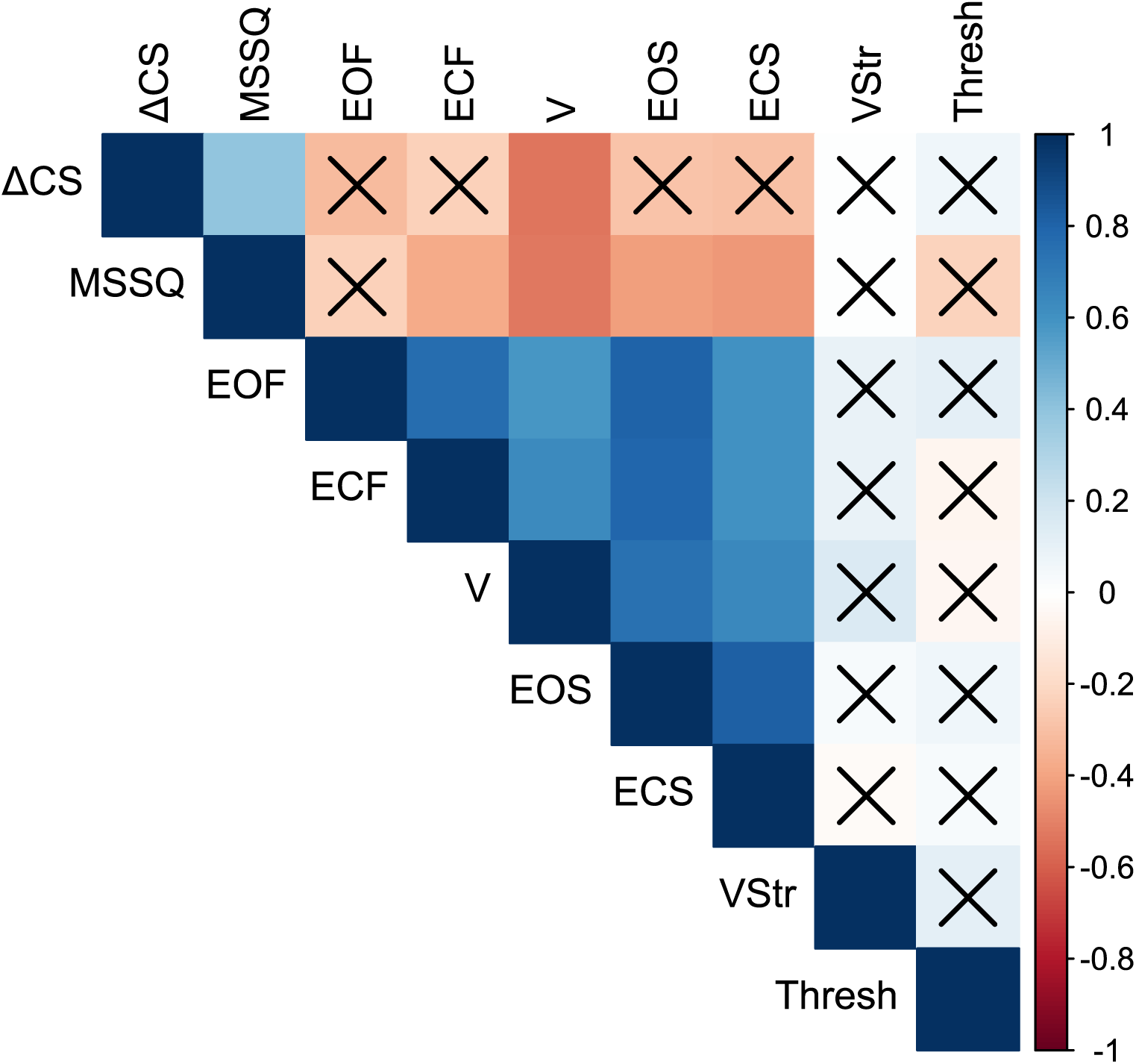
Correlation plot for measures obtained in the study. Correlations, represented by colour, are Pearson *r*(28) values, which are significant (*p* < .05) unless crossed with an X. CS = Cybersickness; MSSQ = Motion Sickness Susceptibility Questionnaire; EOF = Eyes Open Foam condition sway path length (SPL); ECF = Eyes Closed Foam condition SPL; V = Vection condition SPL; EOS = Eyes Open Standard condition SPL; ECS = Eyes Closed Standard condition SPL; VStr = Vection Strength ratings; Thresh = Vestibular Thresholds.

The balance control measures across the five sensory conditions were highly correlated. The Pearson *r* correlation values ranged from 0.59 to 0.82 (average value of 0.70). This suggests that the amount of sway demonstrated by a participant in one condition was predictive of the participant’s balance control in other sensory conditions, consistent with previous literature (Horak, 1987; Winter et al., 2003, 1998).

We observed negative correlations between ∆CS and total sway path length in every balance control condition, with a mean Pearson *r* value of −0.33. Of these conditions, the only significant correlation between sway path length and ∆CS was in the vection condition (*r*(28) = −0.53, *p* = .002). A scatterplot depicting this relationship is shown in Figure 9. We also observed a significant correlation between sway path length in the vection condition and previous history of motion sickness susceptibility (MSSQ; *r*(28) = −0.52, *p* = .003).

**Figure 9.**
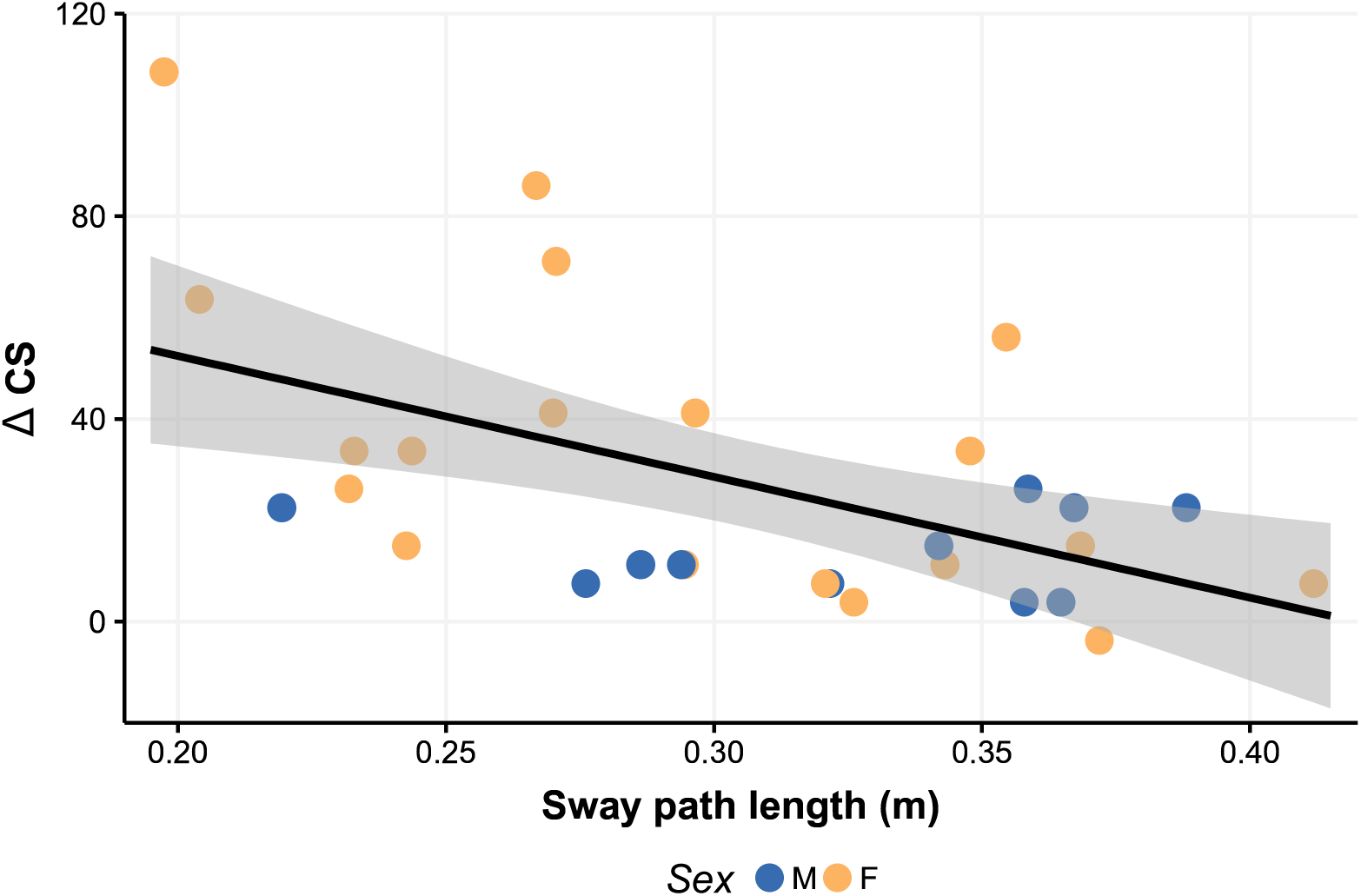
Scatterplot showing the relationship between average sway path length and ∆CS in the V (vection) balance control condition. Participant sex in color (*M* = Male, *F* = Female). Shaded area depicts 95% confidence interval. Note: Although there was one participant with higher ∆CS, it did not constitute a statistical outlier (Dixon one-sided outlier test, *Q* = 0.36, *p* = .07) and removal of this datapoint would still have resulted in a significant negative correlation (*r*(27) = −0.43, *p* = .019).

With respect to participant sex, we found that ∆CS significantly differed between male and female participants (Kruskal-Wallis *X*^2^(1) = 3.95, *p* = .047), with women experiencing more ∆CS (*M* = 34.84, *SEM* = 6.97) than men (*M* = 13.94, *SEM* = 2.48). There was no difference between men and women with respect to MSSQ total scores (Kruskal-Wallis *X*^2^(1) = 1.61, *p* = .204). Additionally we found no differences in total sway path length between men and women in any balance condition (Kruskal-Wallis tests, *X*^2^(1) < 1.84, *p*s ≥ .175).

We obtained several non-significant tests for the effects of behavioural or demographic factors on CS. There was no correlation between ∆CS scores and participant age (*r*(28) = 0.20, *p* = .297), vestibular thresholds (*r*(28) = 0.06, *p* = .743), or average vection strength ratings (*r*(28) = 0.01, *p* = .964). In addition, there was no effect of the self-reported daily activity level of the participant (Kruskal-Wallis *X*^2^(2) = 1.05, *p* = .590), or ethnicity (Kruskal-Wallis *X*^2^(2) = 0.88, *p* = .349) on ∆CS scores.

There were no significant correlations between vestibular thresholds and any other factor (*r*s(28) ≤ 0.22) or between vection strength and other factors (*r*s(28) ≤ 0.16). The strongest correlation for vestibular thresholds was with MSSQ scores (*r*(28) = 0.22, *p* = .23).

### 3.2. Multiple regression analysis

In order to estimate the contribution of each candidate factor to CS, we constructed a multiple regression model. However, our dataset included a set of highly multicollinear variables, such as the sway measures obtained in the five balance control conditions. As well, MSSQ scores were significantly correlated with scores on some balance control conditions (see Figure 8 above). This issue precludes a standard multiple regression approach. Instead we used Principal Components Regression (PCR), which involves conducting a principal components analysis (PCA) on the dataset and then subjecting the principal components to a multiple regression model. PCA is an unsupervised dimensionality reduction technique that results in uncorrelated linear combinations of variables (principal components, PCs) that are ordered by the amount of variance they explain in the original dataset. This procedure eliminates multicollinearity at the expense of the interpretability of the predictors (Massy, 1965).

We identified eight factors to be used in the PCA. These factors were: Motion sickness susceptibility (MSSQ); vestibular thresholds; vection magnitudes; and total sway path length measures from the five balance conditions. In PCA, the first PC always explains the most variance, and in our dataset PC1 explained 50.96% of variance in the original dataset, while PCs 2-4 and PCs 5-8 carried approximately 37% and 12% of the remaining variance, respectively (see Figure 10 for a scree plot of PC variances). While the PCs are less readily interpretable than the original factors, insight into what they represent can be gained by inspection of the factor loadings for each PC. Figure 11 depicts these factor loadings, where higher values indicate a greater expression of that factor in the PC. For instance, it can be determined from Figure 11 that PC1 primarily represents a linear combination of all five balance control conditions and some variance related to MSSQ scores.

**Figure 10.**
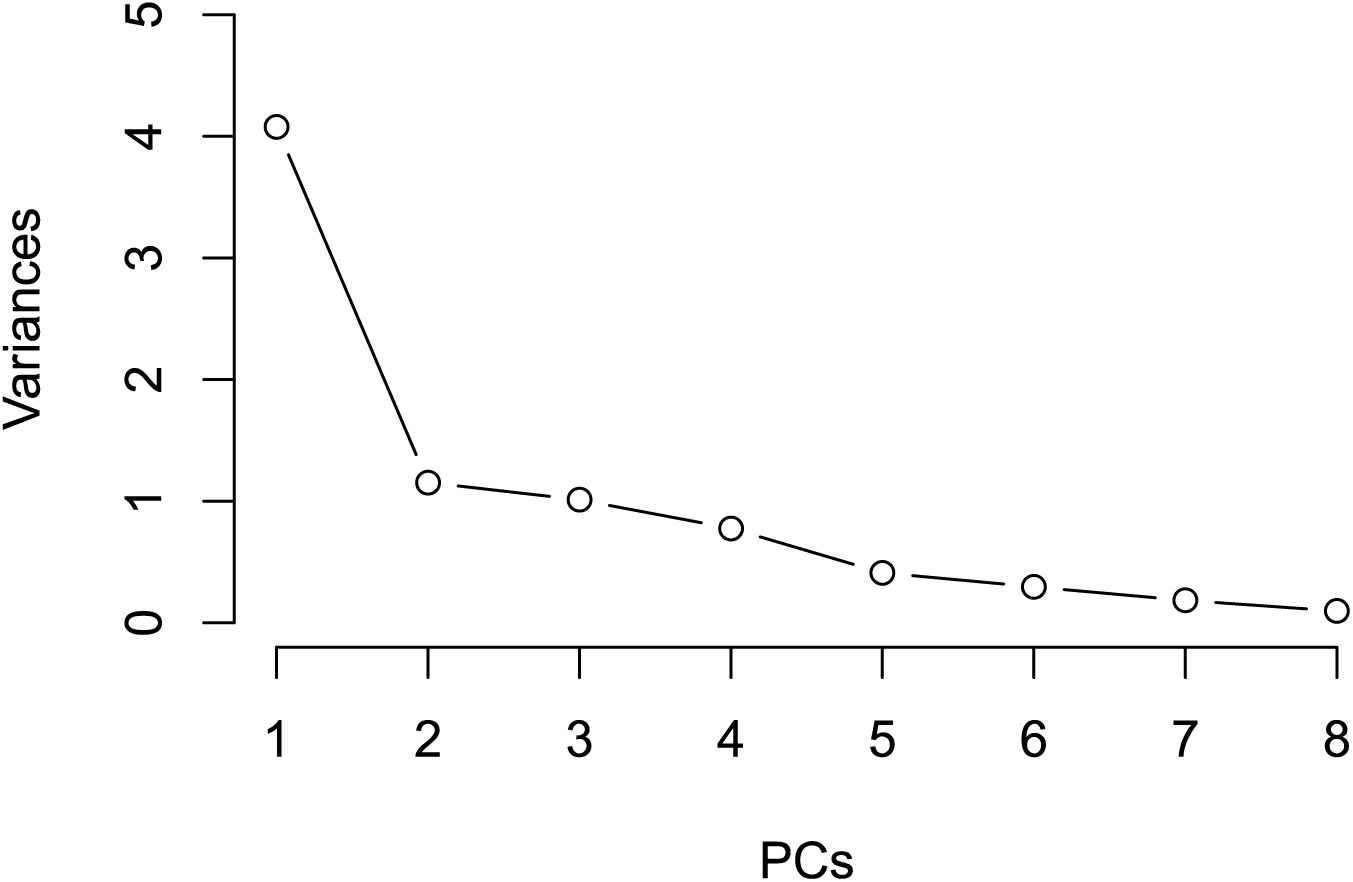
Scree plot showing variances in the initial dataset accounted for by each PC (eigenvalues).

**Figure 11.**
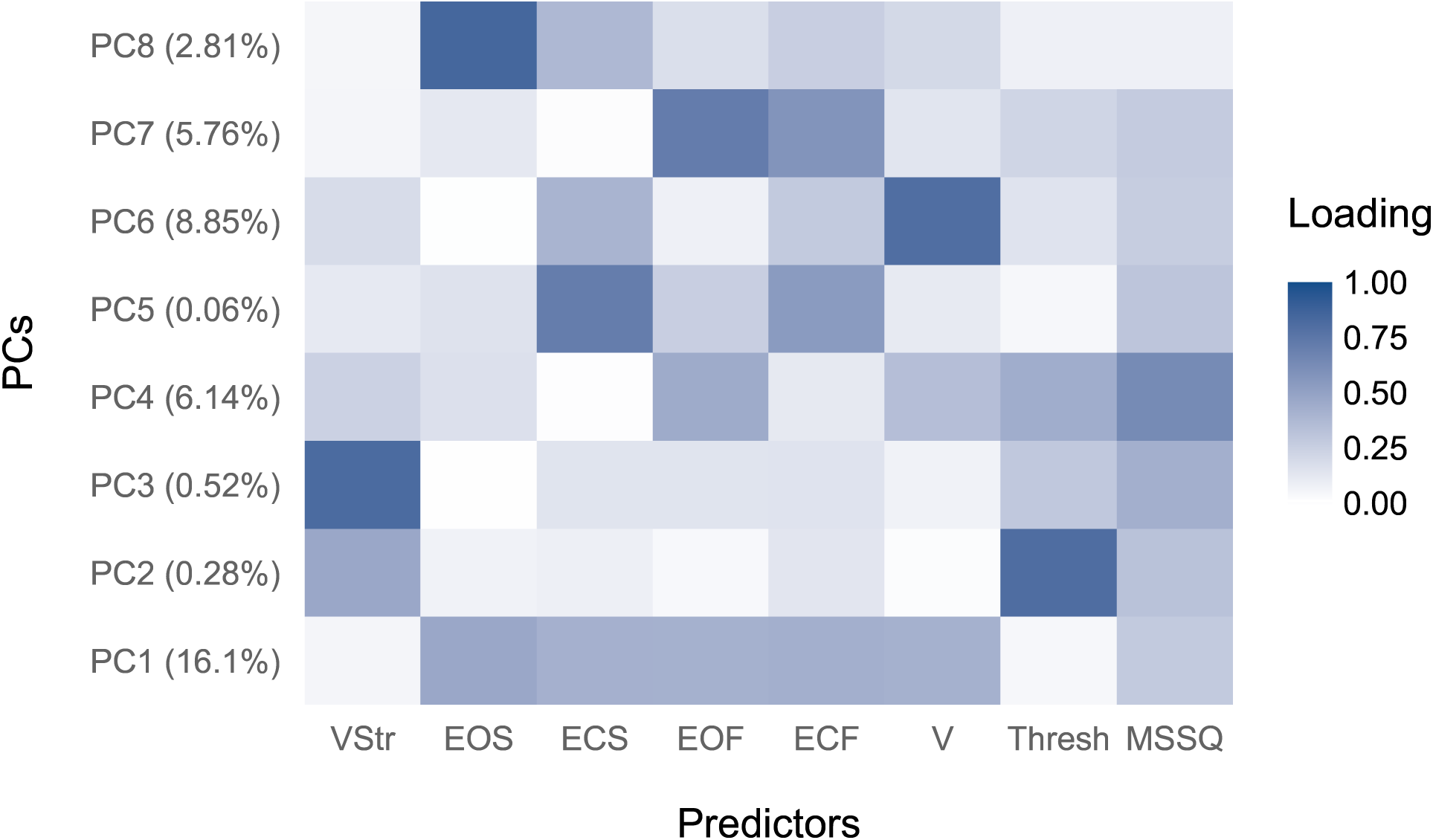
Factor loadings for each PC. Percentage values on left indicate the amount of variance in ∆CS scores accounted for by each PC. Dark blue depicts higher loadings, representing a greater expression of that factor on the PC. Abbreviations are as in Figure 8.

We selected the components to be entered into the regression model based on their correlation with ∆CS scores, with a pre-determined criterion value of *r* = 0.20 (e.g., Dennison et al., 2016; Kim et al., 2005). Predictors were entered into the model simultaneously to avoid the problems of stepwise regression techniques (Stephens et al., 2005). The predictors that met this criterion were: PC1, PC4, PC6, and PC7 (Table 1). The results of the PCR revealed that the combination of these four orthogonal predictors significantly accounted for 36.8% of the variability in ∆CS scores (*R*^2^ = 0.37, *R*^2^*_Adj_* = 0.27, Cohen’s *f*^2^ = 0.59, *F*(4, 25) = 3.64, *p* = .018). While PC6, PC4, and PC7 were not significant predictors alone (*p*s ≥ .07), the first principal component explained 16.1% of the variance in ∆CS scores, and this was revealed to be a significant component of the regression model (*p* = .018, see Table 1).

**Table 1.**
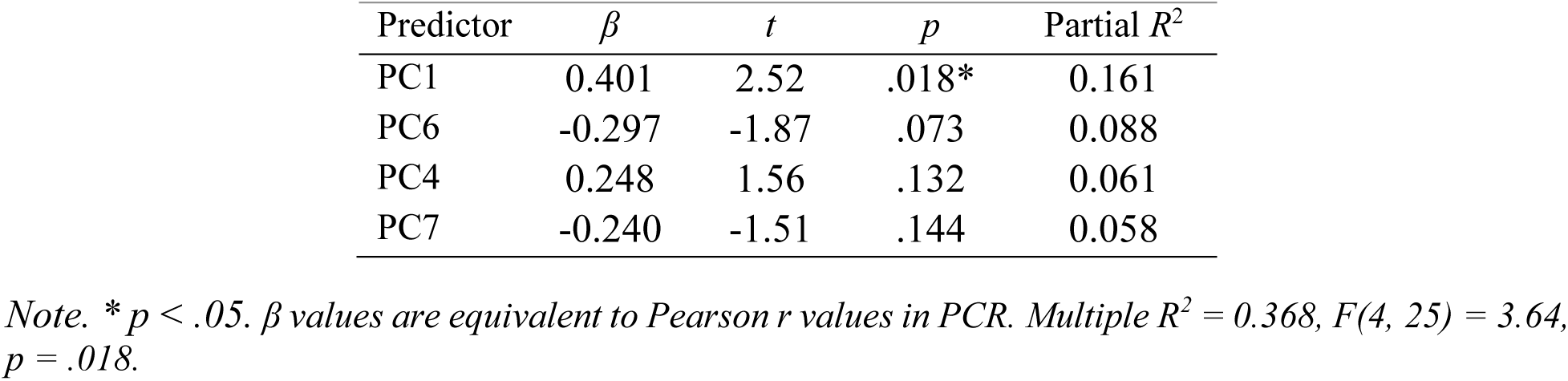
Principal Components Regression Parameters for Predicting CS Scores.

| Predictor | $\beta$ | $t$ | $p$ | Partial $R^2$ |
| --- | --- | --- | --- | --- |
| PC1 | 0.401 | 2.52 | .018* | 0.161 |
| PC6 | -0.297 | -1.87 | .073 | 0.088 |
| PC4 | 0.248 | 1.56 | .132 | 0.061 |
| PC7 | -0.240 | -1.51 | .144 | 0.058 |

*Note.* \* $p < .05$ . $\beta$ values are equivalent to Pearson $r$ values in PCR. Multiple $R^2 = 0.368$ , $F(4, 25) = 3.64$ , $p = .018$ .

Given that PC1 primarily represents a sign-reversed coding of sway values (e.g., see Figure 12A), the sign of beta for PC1 was positive. This reflects the negative correlation between ∆CS and the linear combination of balance control measures. On the other hand, for PC6 (which mainly expresses sway in the vection condition), the PC encodes high sway as positive and vice-versa (Figure 12B). Therefore, the negative beta sign shows a negative relationship between sway and ∆CS. The same can be said for PC7 (primarily foam balance control conditions), where the negative beta sign depicts the inverse relationship between total sway in these conditions and ∆CS scores that we reported previously. While loadings on PC4 are spread more equally across predictors, the strongest loading is for MSSQ scores where the positive sign of beta reflects the positive correlation between MSSQ and ∆CS scores.

**Figure 12.**
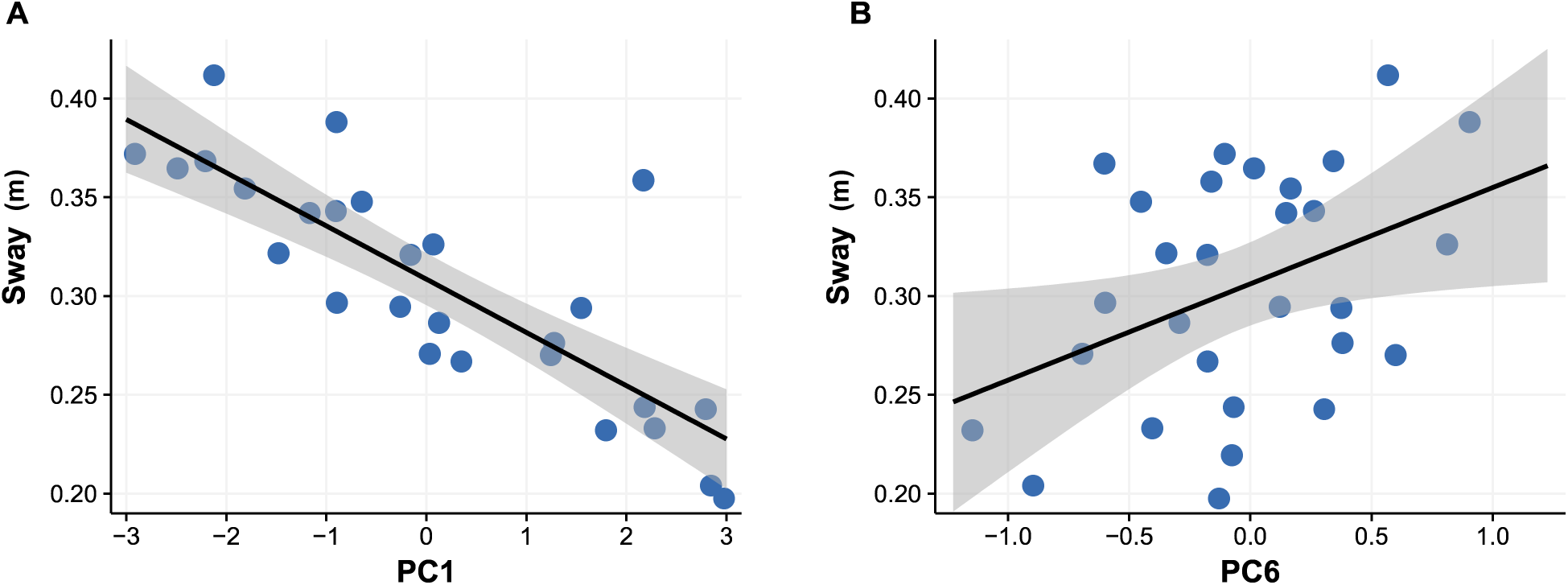
Scatterplot of total sway path length in the vection balance control condition on the ordinate against scores on (A) PC1 or (B) PC6 on the abscissa. Shaded areas are 95% confidence intervals.

### 3.3. Model comparison

We aimed to determine if the model that we selected based on correlation between PCs and CS was the optimal model. An alternative would be that a saturated model (all PCs as predictors) predicted ∆CS significantly z model (PC1 alone) predicted ∆CS equally as well as our chosen model (and would therefore be considered ‘optimal’ due to parsimony). We calculated the Akaike Information Criterion (AIC) for all models, and adopted a delta-AIC value of 2 as a criterion for preferring the lower AIC model. Results are shown below (Table 2). The model based on correlations between PCs (PC1, PC4, PC6, & PC7) exhibited the best model fit.

**Table 2.** Akaike Information Criterion (AIC) for Multiple Regression Models.

| Predictors (PCs) | AIC | delta-AIC |
| --- | --- | --- |
| 1, 4, 6, 7 | 278.97 | . |
| All (saturated) | 285.17 | -6.20 |
| 1 | 281.49 | -2.52 |

## 4. Discussion

Here we aimed to estimate the contributions of several candidates thought to play a role in CS, including vestibular thresholds, balance control, and vection susceptibility. Correlation analysis revealed a significant negative relationship between ∆CS and sway path length when participants observed a vection stimulus. We conducted a PCR to assess the contribution of orthogonal linear combinations of each candidate factor on ∆CS and interpreted the factor loadings for the significant predictors. We found strong evidence supporting the role of vection susceptibility and the role of balance control in CS. The correlation between balance control and ∆CS that we observed was negative, opposite to that which was reported in previous literature (e.g., Chardonnet et al., 2017; Stoffregen & Smart, 1998; Takada et al., 2007), although a negative correlation has also been identified in other recent work (Dennison & D’Zmura, 2017, 2018; Sadiq et al., 2017). We found no evidence of a link between ∆CS and vestibular thresholds or verbal ratings of vection strength. Our results demonstrate that behavioural and self-report data gathered before exposure to VR can be used to assess the likelihood of CS emerging on an individual basis.

### 4.1. Comparison with previous regression studies

Previous work has adopted a regression approach to CS prediction. Kim and colleagues (2005) ran a stepwise regression analysis using several predictors including MSSQ scores and physiological data. A combination of electrocardiography (ECG), electroencephalography (EEG), and MSSQ scores predicted 46% of variability in CS. However, the model of Kim and colleagues was comprised mostly of measurements taken during VR exposure (with the goal of CS classification), whereas each of the measures obtained in the current study were from pre-exposure assessments (with the goal of CS prediction). It is therefore not surprising that the variance explained by our model is lower than slightly lower than 46%. Indeed, the accuracy of our model is almost identical to a classification model produced by Dennison and coworkers (2016) who found that online measurements (during VR exposure) of bradygastric power, breathing rate, and blink rate obtained during VR exposure predicted 38% of variance in CS scores. In the current experiment our aim was to predict ∆CS using data that we obtained prior to VR exposure, but we anticipate that combining pre-test and on-line physiological data would account for an even larger proportion of variance in CS.

Although the regression approach has produced successful findings, other computational techniques may prove more powerful with respect to prediction. While a recent study reported high classification accuracy using linear discriminant analysis (LDA) with physiological data to categorize whether participants were viewing visual stimuli on a monitor or with a VR display (Dennison et al., 2016), future efforts should employ techniques such as LDA or support vector machines (SVM) to construct predictive models of CS derived from behavioural and physiological measures.

### 4.2. Balance control

The theory that postural instability precedes motion sickness (Riccio & Stoffregen, 1991) has proven extremely influential in research on CS, and the predictions of the theory have been supported several times (e.g., Chardonnet et al., 2017; Stoffregen & Smart, 1998; Takada et al., 2007). Our results suggest that individual differences in balance control can predict tolerance of nauseogenic stimuli, but the direction of the relationship we identified between sway path length and ∆CS was negative. Our results agree with the findings of a separate body of literature wherein a null or negative association between sway and CS was found (e.g., Dennison & D’Zmura, 2017, 2018; Sadiq et al., 2017). For instance, it has been revealed that postural instability caused by visual perturbations does not produce CS as predicted by the postural instability account of motion sickness (Dennison & D’Zmura, 2018). Dennison and D’Zmura (2017) also showed that participants who exhibited less postural sway than others were more likely to experience CS. Those authors concluded that participants tended to reduce their body motion if they experienced CS, implicating CS as the cause of lower body sway, rather than vice-versa. Our current results rule out this ‘VR lock’ explanation given that we measured balance control before exposure to VR content, and obtained negative correlations in all five conditions (although most trends were non-significant). However, we contend that our results do not directly contradict the postural instability theory of Riccio and Stoffregen (1991). Dynamic stability of posture does not necessitate a rigidly stationary body and low sway path length (Błaszczyk, 2008; Cho et al., 2014; Palmisano et al., 2018). Increased postural sway could demonstrate a more flexible balance control system and a readiness to adapt to novel sensorimotor conditions such as those presented by VR. A control strategy where the individual struggles to avoid postural adjustments in response to a compelling visual stimulus may minimize body sway. At the same time, this may constitute precisely the type of ineffective strategy that Riccio and Stoffregen (1991) identify as a precursor to motion sickness. As such, the current results complement the idea that balance control measures are valuable predictors of susceptibility to CS, although whether high-CS participants were ‘more stable’ or ‘less stable’ than others is open to interpretation.

While it is possible that the difference between our results and other research can be attributed to methodological differences, the method we used to record balance control (30 seconds of standing in quiet stance prior to VR exposure) was similar to other research. For instance, Chardonnet and coworkers (2017) recorded balance for 30 seconds pre- and post-exposure to VR, and used these data to calculate sway area. Palmisano and colleagues (2018) measured vection for 30 seconds and, in a separate task, they measured balance control during quiet stance for 60 seconds. The balance data were used to conduct recurrence quantification analysis, a non-linear measure for balance control. Other studies are unclear with respect to trial durations (e.g., Palmisano et al., 2017). In addition, although our measure of CS (∆CS) is different to most other studies, due to the high correlation between ∆CS and SSQ scores for the nauseogenic VR content used in our experiment (*r* = .95), we do not consider it likely that this accounts for the difference in results.

### 4.3. Vection

Although some have proposed that vection plays a strong role in motion sickness (Keshavarz et al., 2015), some previous research has been unable to identify a strong relationship between vection and visually-induced motion sickness (Palmisano et al., 2018). Our results extend this research by showing that vection responses predict CS in a nauseogenic virtual environment. We found that the sway in response to vection stimuli had the strongest predictive power for CS among all measures collected here, being that these data strongly loaded onto the two most important PCs in the regression model (1 and 6). This result presents the possibility of using sway responses to vection as a simple predictive tool for individual susceptibility to CS while avoiding nauseogenic conditions entirely (we note however that similar vection stimuli can produce motion sickness in wide field-of-view conditions; Palmisano et al., 2018, 2017).

While others have identified a positive association between vection strength and motion sickness (Hettinger et al., 1990; Hettinger & Riccio, 1992), our results complement other work that has identified a negative relationship between CS and vection susceptibility as measured by magnitude ratings (Palmisano et al., 2017). Although these authors suggested that the negative relationship they found was an artefact of their experimental design, our results constitute a replication of this effect in different settings, suggesting the relationship may be a reliable one.

Our results show that ratings of vection strength neither correlated with sway path length, nor did they provide a strong predictive value for CS ratings. This suggests a dissociation between the behavioural and subjective components of the illusion. While other studies have found a correlation between these measures, the association depends on the visual display conditions with respect to factors such as visual eccentricity and foreground vs. background interpretation (Delorme & Martin, 1986; Kawakita et al., 2000). Importantly, Delorme and Martin (1986) highlight the frequent occurrence of cases where participants had synchronous postural reactions to optic flow stimulation without reporting any subjective vection responses. Our choice of an oscillating vection stimulus may have reduced the relationship between sway and vection strength: Palmisano and colleagues (2014) documented that the relationship between vection responses and postural sway was strong for smooth-vection stimuli, but weak for jittering or oscillating vection stimuli. Another point of consideration is that participants in the present study did not experience very strong vection overall based on subjective strength ratings. However, there was significantly greater sway path length observed in the vection condition compared to the eyes-open standard condition, reflecting that the optic flow stimulus used was sufficient for inducing vection.

### 4.4. Vestibular sensitivity

The vestibular threshold data we obtained were broadly in line with the results of others who have assessed the thresholds for small yaw rotations at similar frequencies (Grabherr et al., 2008). The most common thresholds in our experiment fell in the range between 0.5-0.7 deg/s, which closely aligns with Grabherr and colleagues’ (2008) average threshold of 0.64 deg/s at 1 Hz. On the other hand, the thresholds we obtained were lower than those measured by Benson and colleagues (1989) who identified ~1 deg/s thresholds at 1.1 Hz rotation, and others reviewed by Soyka and colleagues (2012), although the difference may be attributed to metholodigical variations between the experiments. Notably, we did not find any relationship between vestibular thresholds and CS as predicted by research on vestibular dysfunction patients (Cheung et al., 1991, 1989; Johnson et al., 1999) and vestibular migraine patients (Money & Cheung, 1983). However, there was a non-significant tendency for vestibular thresholds and past history of motion sickness to be correlated, in line with previous research showing a link between increased motion sickness in individuals with high vestibular sensitivity (Kuritzky et al., 1981). Here our vestibular threshold measurements were limited to the yaw axis due to experiment duration constraints. It may be that a more comprehensive measurement of vestibular thresholds (e.g., a ‘vestibulogram’, Grabherr et al., 2008) would help to clarify if the prediction of CS can be achieved using vestibular sensitivity to rotation in other axes or at other frequencies to those tested here.

### 4.5. Other candidate factors

Notably, while our regression model significantly predicted CS, there remained approximately 63% of variability in CS that was unexplained by the combination of our predictors. There are several candidates related to sensorimotor processing that merit exploration in the search for higher variance-explained. These include genetic factors, the contribution of which should be assessed with a large sample. Several genes have recently been identified as linked to motion sickness in a full genome sequencing study (Hromatka et al., 2015), but a genome-wide approach to CS susceptibility would not be possible barring a large proportion of the population undertaking a standardised VR experience. However, it appears likely that inherited factors are a latent source of multicollinearity between the sensorimotor indices measured in our experiment, and future high-power studies will be needed in order to investigate this possibility. Other promising latent factors include cue weightings across modalities, which may modulate the likelihood of detecting artifacts in the sensory streams. The relationship between CS and an individual’s ability to rapidly reweight multisensory cues in conditions of mismatch is also understudied (for a recent review see Gallagher & Ferrè, 2018). While some authors have assessed the speed of multisensory reweighting and linked this to carsickness (Balter et al., 2004; note, no relationship was observed), there is a need to examine the reweighting function in VR conditions. In addition, an individual’s tendency to bind near-synchronous multisensory cues (the ‘temporal binding window’, Dixon & Spitz, 1980; Wallace & Stevenson, 2014) may predict the likelihood of sickness emerging in environments where a barrage of multimodal cues with varying latencies must be processed by the central nervous system.

### 4.6. Implications for rehabilitation

The near future of motor rehabilitation initiatives involves a substantial use of VR technology (Guo et al., 2017; Kim et al., 2018). The results obtained here illustrate the complex factors that may determine if VR is well-tolerated or not, including several sensorimotor processing indices such as balance control. The sudden changes in the body control strategy that are imposed by an injury could produce maladaptive responses to multisensory information (Riccio & Stoffregen, 1991). Do individuals with temporarily reduced balance control experience increased CS? The current research reinforces the likelihood that an individual’s balance control plays some part in the degree to which they experience CS, but we are unable to answer if sudden impairments to the sensorimotor control system are associated with CS. This question is of significant current interest given the challenge of performing skills training or rehabilitation in nauseating conditions.

## 5. Conclusion

Here we have shown that a combination of factors measured prior to experiencing intense VR content holds significant predictive power for the amount of CS an individual experiences. The results indicate that the more a participant swayed in response to vection stimuli, the less CS they were likely to experience. Although our predictive model for CS depicts a central role for the destabilising effect of vection on postural stability, we found utility in other measures, such as balance control while standing on foam, and self-reports of motion sickness susceptibility. These results are intended to guide the development of future efforts to predict CS prior to its experience, which we expect to require multifactorial data collection and dimensionality reduction techniques such as those employed here. The knowledge and understanding gained from this research about the links between self-motion perception, balance control, and CS may also prove useful in other domains, such as guiding balance rehabilitation initiatives.

## Acknowledgements

We are grateful to Jeff Rice for technical assistance in this project. This research was supported by grants to MBC from Oculus Research, the Ontario Research Fund and Canadian Foundation for Innovation’s John R. Evans Leaders Fund (#32618), and the Natural Sciences and Engineering Research Council of Canada (#RGPIN-05435-2014).

